# Cryo-EM of dynein microtubule-binding domains shows how an axonemal dynein distorts the microtubule

**DOI:** 10.1101/590786

**Authors:** Samuel E. Lacey, Shaoda He, Sjors H. W. Scheres, Andrew P. Carter

## Abstract

Dyneins are motor proteins responsible for transport in the cytoplasm and the beating of the axoneme in cilia and flagella. They bind and release microtubules via a compact microtubule-binding domain (MTBD) at the end of a long coiled-coil stalk. Here we address how cytoplasmic and axonemal dynein MTBDs bind microtubules at near atomic resolution. We decorated microtubules with MTBDs of cytoplasmic dynein-1 and axonemal dynein DNAH7 and determined their cryo-EM structures using the stand-alone Relion package. We show the majority of the MTBD is remarkably rigid upon binding, with the transition to the high affinity state controlled by the movement of a single helix at the MTBD interface. In addition DNAH7 contains an 18-residue insertion, found in many axonemal dyneins, that reaches over and contacts the adjacent protofilament. Unexpectedly we observe that DNAH7, but not dynein-1, induces large distortions in the microtubule cross-sectional curvature. This raises the possibility that dynein coordination in axonemes is mediated via conformational changes in the microtubule.

## Introduction

The dynein family is a group of minus-end directed microtubule motors. The two cytoplasmic dyneins (dynein-1 and dynein-2) are involved in long-range movement of cellular cargoes (Reck-Peterson et al., 2018; Roberts et al., 2013). Multiple inner and outer arm axonemal dyneins power the beating motion in cilia and flagella by sliding adjacent doublet microtubules past each other (Satir et al., 2014). All dynein family members share a common architecture, based around a heavy chain that contains a cargo-binding tail region and a force-generating motor domain. The motor consists of a ring of six connected AAA+ subdomains (AAA1-6) with the nucleotide cycle of the first, AAA1, powering movement (Schmidt and Carter, 2016). Dyneins bind to microtubules via a small microtubule-binding domain (MTBD) consisting of 6 short helices (H1-H6) (**Figure 1A**). The MTBD is connected to the AAA+ ring by an antiparallel, coiled-coil stalk, containing helices CC1 and CC2. It binds to the microtubule at the tubulin intradimer interface (Carter et al., 2008).

**Figure 1.**
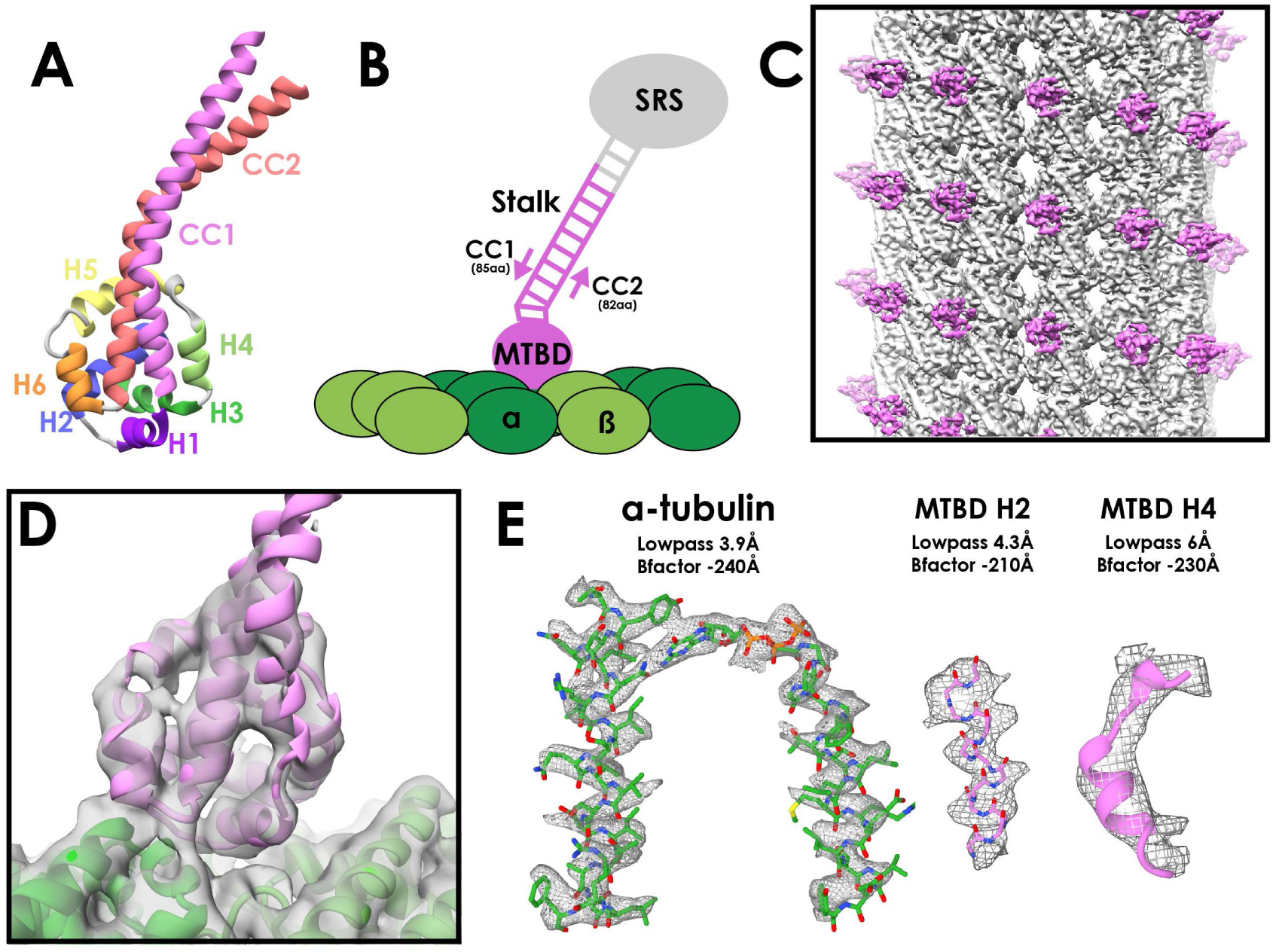
**A** – Crystal structure of the cytoplasmic dynein-1 MTBD in the low-affinity β+ registry (PDB 3ERR) colored by helix. **B** – Schematic of the MTBD constructs used for structure determination. A globular seryl-tRNA synthetase (SRS, grey) has a protruding coiled-coil to which 12-heptads of the dynein stalk is fused (pink). CC1 is 3 residues longer than CC2 to force the stalk into the high-affinity α registry, allowing the MTBD to bind to the microtubule (green) **C** – Reconstruction of the cytoplasmic dynein-1 MTBD (pink) bound to microtubule (grey), lowpass-filtered to 5Å. **D** – New models for the cytoplasmic dynein MTBD (pink) and tubulin (α-tubulin dark green, β-tubulin light green) was refined into the cryo-EM density (lowpass-filtered to 5Å) **E** – Representative density of different regions of the map, filtered and sharpened according to local resolution.

Nucleotide dependent conformational changes are transmitted to the MTBD through the stalk to modulate its affinity for microtubules, allowing dynein to bind and release as it steps along the microtubule. The stalk transmits these conformational changes due to its ability to pack into two stable registries (α and β+), representing a half-heptad shift of CC1 relative to CC2 (Carter et al., 2008; Gibbons et al., 2005; Kon et al., 2009, 2012; Schmidt et al., 2015). The structure of the MTBD in the low microtubule affinity state (stalk in β+ registry) has been solved to high-resolution (Carter et al., 2008; Nishikawa et al., 2016). The relative movement of CC1 towards the ring (stalk in α registry) pulls on the MTBD to create a high affinity state. The structure of this state was visualized in a landmark 9.7Å cryo-electron microscopy (cryo-EM) structure of the MTBD bound to microtubules (Redwine et al., 2012). A pseudoatomic model was fit into the cryo-EM map using molecular dynamic simulations, and showed large conformational changes throughout the MTBD upon binding. The authors concluded that specific interactions between the microtubule and H1, H3 and H6 of the MTBD are required to induce and maintain this high-affinity conformation.

A high degree of conservation in the MTBD between the different dynein family members suggests that this mechanism of microtubule binding is universal (Hook and Vallee, 2006). However, many axonemal dyneins have an insertion between H2 and H3 called the flap, the function of which is unclear. An NMR structure of the *Chlamydomonas reinhardtii* flagellar dynein-c MTBD showed that the flap consists of two flexible beta-strands extending from the MTBD core (Kato et al., 2014). The flap was predicted to sterically clash with the microtubule surface, and therefore undergo rearrangement upon binding.

We decided to take advantage of recent technological advances in cryo-EM (Fernandez-Leiro and Scheres, 2016; He and Scheres, 2017) to compare the structures of a cytoplasmic and axonemal dynein MTBDs bound to microtubules. We determined the structure of mouse cytoplasmic dynein-1 MTBD on microtubules to an overall resolution of 4.1Å. We observe a number of structural differences to the 9.7Å structure. This leads to an updated model for the transition from a low- to high-affinity state, based only around movement of H1 to avoid steric clashes with the microtubule surface. Furthermore, we determine the structure of microtubules decorated with the MTBD of the human inner-arm dynein DNAH7 to 4.8Å resolution. We show that its flap contacts an adjacent protofilament from the rest of the MTBD, and show that this interaction dramatically distorts the microtubule cross-sectional shape.

## Results

### Structural determination of cytoplasmic dynein-1 MTBD decorating microtubules

To fix the stalk in the high microtubule affinity α-registry we used a chimeric fusion construct (SRS-DYNC1H1_3260-3427_) in which the mouse cytoplasmic dynein-1 MTBD and 12 heptads of stalk are fused to a seryl-tRNA synthetase (**Figure 1B**) (Carter et al., 2008). Predominantly 13-protofilament microtubules were made by polymerizing tubulin in a MES-based buffer (Pierson et al., 1978). The MTBD and MT were incubated together on-grid and vitreously frozen for cryo-EM (**Figure S1A, Table 1**).

**Table 1.** Data collection statistics for the presented structures. * 13-fold non-crystallographic symmetry applied

|  | **SRS-DYNC1H1**<sub>3260-3427</sub> | **DYNC1H1**<sub>1230-4646</sub> | **SRS<sup>+</sup>-DNAH7**<sub>2758-2896</sub> |
| --- | --- | --- | --- |
| Microscope | Krios | Polara | Krios |
| Detector | Falcon III | Falcon III | Falcon III |
| Voltage (kV) | 300 | 300 | 300 |
| Exposure time (s) | 1.5 | 1.5 | 1.5 |
| Total dose (e<sup>-</sup>/Å<sup>2</sup>) | 60 | 67.5 | 55.5 |
| Pixel Size (Å) | 1.04 | 1.34 | 1.1 |
| Defocus Range (um) | -1.5 to -4.5 | -1.5 to -4.5 | -1.5 to -4.5 |
| Sessions | 1 | 1 | 1 |
| Micrographs | 1995 | 2455 | 1995 |
| Symmetry | 13* | 13* | 13* |
| Final particles | 59100 | 38142 | 41984 |
| Resolution (Å) | 4.1 | 5.5 | 4.8 |

Microtubules are characterized by having like-for-like lateral contacts (i.e. α-to-α tubulin contacts), however most microtubule architectures, including 13-protofilament microtubules, break this pattern with a seam of heterotypic α-to-β contacts (Desai and Mitchison, 1997) (**Figure S1B**). As such, these microtubules cannot be subjected to conventional helical symmetry averaging. Previously, EM structures of microtubules with a seam have used iterative 2D cross-correlation to synthetic projections followed by 3D refinements in order to locate the position of the seam (Sindelar and Downing, 2007; Zhang and Nogales, 2015). We propose an alternative image processing approach that is integrated into the helical Relion pipeline (**Figure S1C**, (He and Scheres, 2017)). 3D refinement follows the standard regime for a C1 helix, however 13-fold local, non-point-group symmetry is applied to the reconstruction following each iteration. The application of local symmetry depends on a user-specified mask and symmetry operators that superimpose equivalent tubulin dimers on top of each other. The application of local symmetry increases the signal in the reference during refinement, and prevents progressive deterioration in the definition of the seam from poorly aligned particles. The same local symmetry is also applied to increase the signal in the final reconstruction. Relion has the advantage of requiring minimal user input, and is capable of sorting sample heterogeneity with 2D and 3D classification.

We initially used 2D classification to remove microtubules that were poorly decorated or possess identifiable non-13 protofilament architectures (**Figure S1C**). Particles from good 2D classes were used for 3D classification, which resulted in a single good class (**Figure S1C**). Manual inspection confirmed that this class contained only 13-protofilament microtubules.

The seam is well defined in the resulting asymmetric reconstruction, with α-tubulins making lateral contact with β-tubulins. This is displayed most clearly when viewing the extended luminal S9-S10 loop in α-tubulin and the additional rise between MTBDs across the seam (**Figure S1E/F**). Application of local symmetry resulted in a map with an overall resolution of 4.1Å (**Figure 1C, 1D, S1D**). Local resolution assessment shows that the tubulin core extends from 3.6Å to 4.4Å resolution (**Figure 1E, S2A**). The tubulin model refined into our density closely matched a previous high-resolution EM model of taxol-stabilised microtubules (Kellogg et al., 2017). The density for the dynein MTBD ranges from 4.4Å to 8Å in resolution (**Figure S2A**). Regions of the MTBD close to the microtubule were of sufficient resolution to see the pitch of alpha helices (**Figure 1E**). We built and refined a new model of the high-affinity form of the MTBD using the low-affinity MTBD crystal structure (Carter et al., 2008) as a starting point (**Figure 1D/E, S2B, Table 2**).

**Table 2.** Model refinement statistics for the presented structures

|  | **SRS-DYNC1H1**<sub>3260-3427</sub> | **SRS<sup>+</sup>-DNAH7**<sub>2758-2896</sub> |
| --- | --- | --- |
| **Map** |  |  |
| Map resolution | 5 | 5.4 |
| Map sharpening B factor | -220 | -200 |
| Map CC | 0.851 | 0.77 |
| Map:Model FSC<sub>0.5</sub> (Å) | 5.11 | 5.54 |
| **Model Composition** |  |  |
| Non-hydrogen atoms | 8210 | 15061 |
| Protein residues | 1035 | 1901 |
| Ligands (GTP/GDP) | 1/1 | 1/1 |
| **R.m.s deviations** |  |  |
| Bond lengths | 0.006 | 0.005 |
| Bond angles | 1.015 | 0.932 |
| **Validation** |  |  |
| MolProbity score | 2.09 | 2.49 |
| Clashscore | 13.61 | 9.83 |
| Poor rotatmers | 0.56% | 6.65% |
| **Ramachandran plot** |  |  |
| Favored (%) | 93.09 | 95.02 |
| Disallowed (%) | 0 | 0 |
| CB deviations (%) | 0 | 0 |

### Microtubule binding involves movement of only one helix

Surprisingly, when comparing the low-affinity crystal structure (Carter et al., 2008) to our new high-affinity structure, we see that the majority of the MTBD undergoes remarkably few changes upon microtubule binding (**Figure 2A/B**). The Cα RMSD between the two structures for helices H2 to H6 is 1.9Å. The major movement involves H1 and CC1, which move together to occupy the intra-dimer cleft above α- and β-tubulin. A more minor change is seen in H6 in order to accommodate its interaction with α-tubulin (**Figure 2A**).

**Figure 2.**
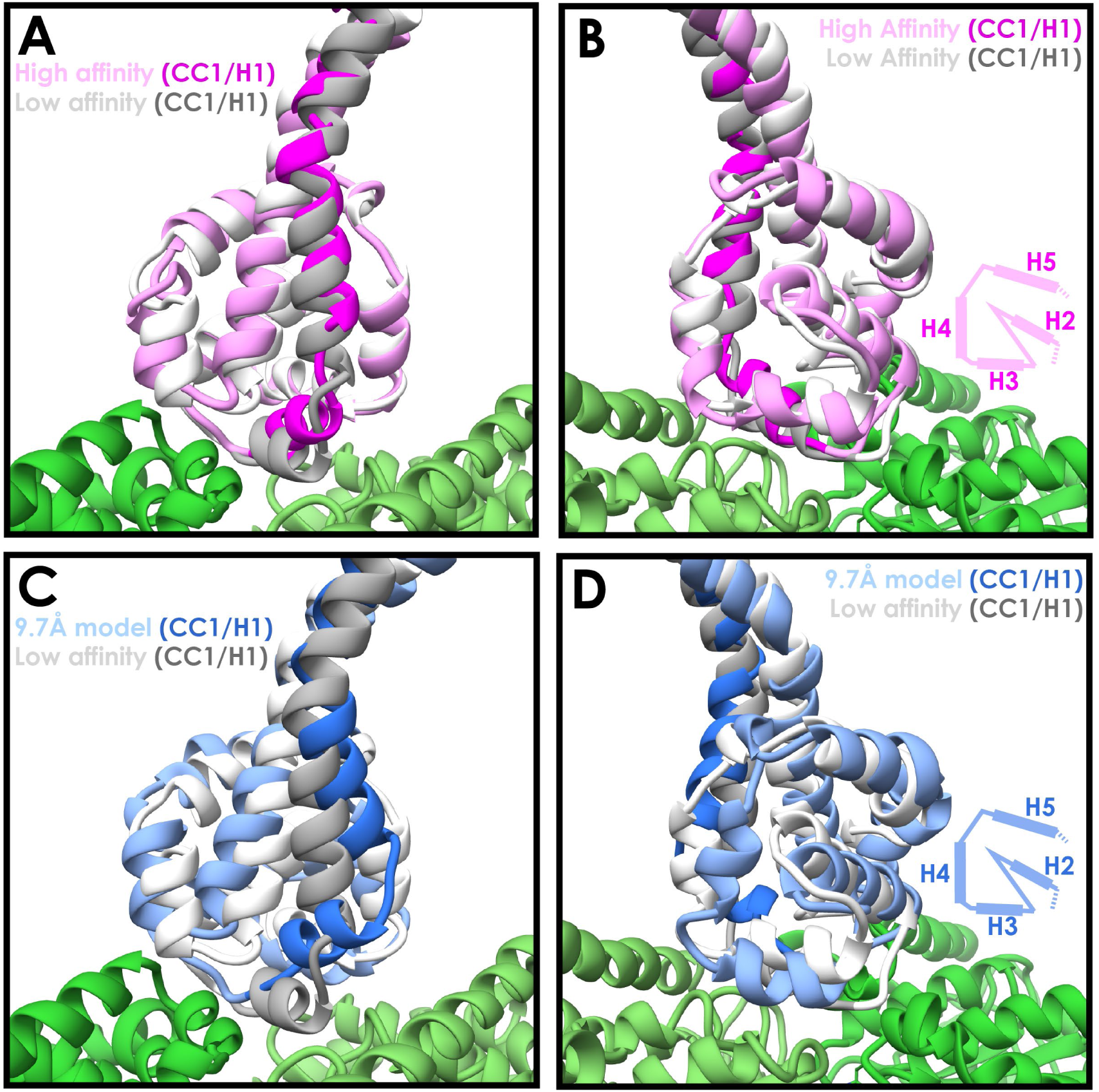
**A** – A comparison between the newly refined cytoplasmic dynein-1 MTBD model (pink) and the low-affinity state crystal structure (PDB 3ERR, docked to the same density, white). CC1 and H1 (highlighted in magenta and grey for high- and low-affinity states respectively) rise in the high affinity state to solve a steric clash with the microtubule (green) **B** – Orthogonal view of A, highlighting the similarity between the two models away from CC1/H1. Cartoon displays organization of H2-H5 as visibile in the model. **C** – A comparison the previous 9.7Å cryo-EM microtubule bound model (blue, PDB 3J1T) and the low-affinity state crystal structure (white, 3ERR). There is a larger movement up and to the side in CC1 and H1 (blue and grey for 3J1t and 3ERR respectively) in 3J1T compared to the new model. **D** – Orthogonal view of C, showing H2, H3 and H4 all in different conformations relative to the microtubule in 9.7Å model.

This is in contrast to the previous 9.7Å microtubule bound model, in which H1 moves towards β-tubulin and shifts H2, H3 and H4 as a result (**Figure 2C/D**, Cα RMSD H2-H6 3.7Å). We see a much smaller movement in CC1 and H1, and as a result H3 and H4 stay much higher above the microtubule surface. The experimental setup used in this study is slightly different to the previous structure. We imaged a longer 12-heptad stalk-SRS construct on 13 protofilament microtubules, compared to a 3-heptad construct on 14-protofilament microtubules. To confirm that the structure we observed is representative of full-length dynein bound to microtubules we imaged a human cytoplasmic dynein-1 motor domain construct (DYNC1H1_1230-4646_, (Steinman et al., 2017) bound to microtubules (**Figure S1A**). Grids were frozen in the absence of nucleotide to attain a high-affinity state. Using the Relion pipeline we achieved an overall resolution of 5.5Å (**Figure S4A, S5A, Table 1**). Lower microtubule occupancy was observed for DYNC1H1_1230-4646_ compared to SRS-DYNC1H1_3260-3427_, which is likely a result of steric clashes between adjacent motor domains around the microtubule. Microtubules with high occupancy were selected following 2D classification. The lower occupancy meant that the MTBD density was at a much lower resolution than the microtubule, and therefore the map was lowpass filtered to 8Å for interpretation. The resulting map fits well with our new model, but H1, H2, H3 and H4 from the 9.7Å model are all at least partially outside the density (**Figure S5C-F**). Accordingly, model-to-map FSC measurements indicate that our new model is a better fit to the map (FSC_0.5_ = 8.66Å (New model) and 12.76Å (9.7Å model) (**Figure S5B**). As such, we conclude that our 4.1Å structure corresponds to the native state of the dynein MTBD bound to microtubules.

The MTBD residues forming the interface with the microtubule in our new model are consistent with previous structural and mutagenesis data (Gibbons et al., 2005; Koonce et al., 2000; Redwine et al., 2012). In contrast, some of the residues on the microtubule previously predicted to interact with the MTBD (Redwine et al., 2012) are too distant in our model (Red bonds, **Figure S5G/H**). For example, K3299 is now over 7Å away from its previously predicted interaction partner β-tubulin E420, but is directly proximal to D427 (**Figure S5G**). Overall, our new model suggests that the MTBD is permanently primed for microtubule binding. The only significant conformational change that occurs upon binding is the movement of H1 upwards to accommodate the change in stalk registry.

### An axonemal dynein MTBD contacts 4 tubulin subunits at once

We next investigated whether the microtubule interaction we observe is conserved across different dynein families. DNAH7 is a monomeric inner arm axonemal dynein closely related to *Chlamydomonas reinhardti* flagellar dynein-c (Hom et al., 2011; Wickstead and Gull, 2007). It is one of 5 human axonemal dyneins to contain a flap insert between H2 and H3 of the MTBD (**Figure 3A + S6A/B**), and has one of the most divergent MTBD sequences compared to cytoplasmic dynein-1.

**Figure 3.**
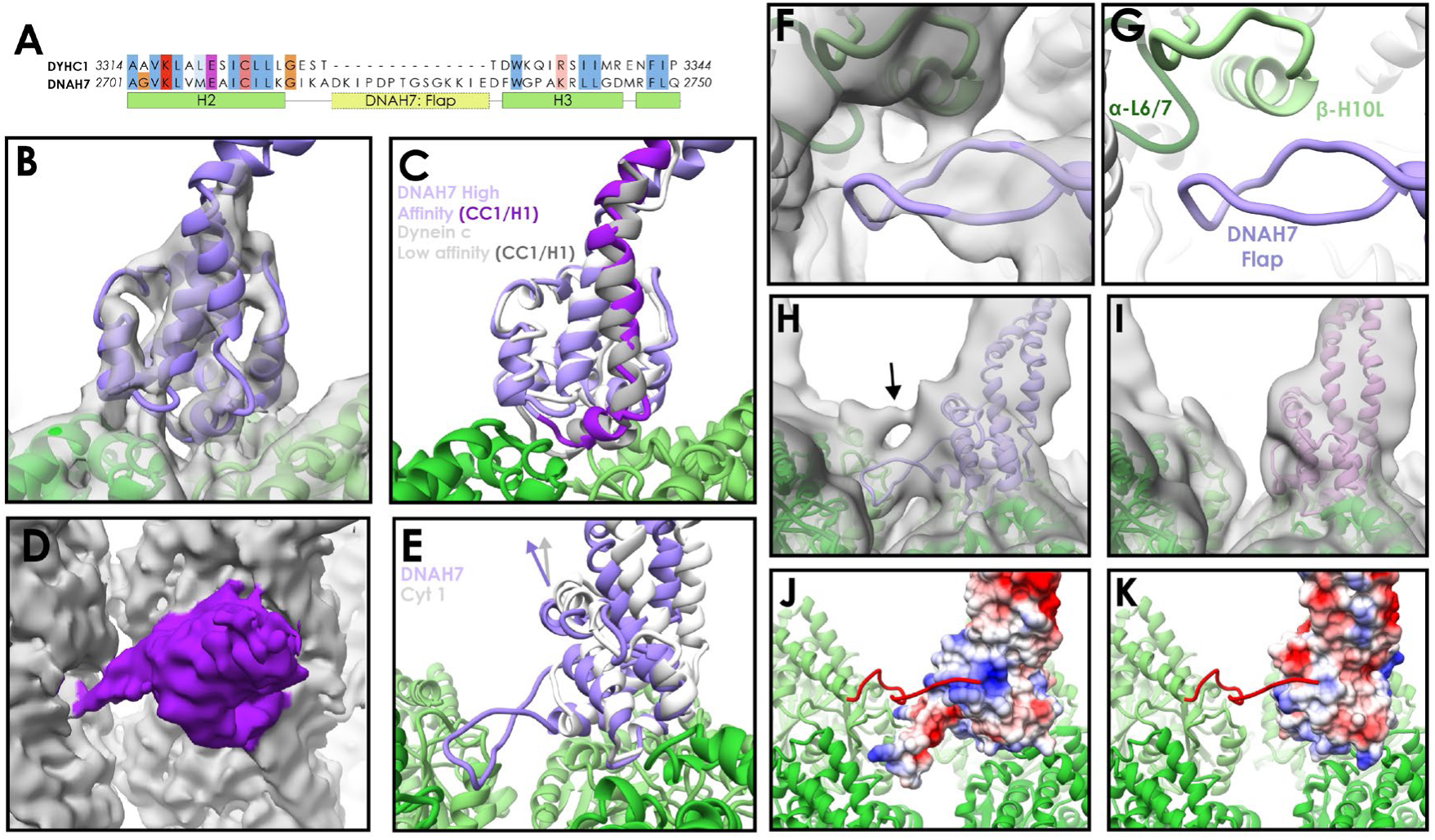
**A** – Partial sequence alignment of human cytoplasmic dynein-1 (DYHC1) and human Axonemal Dynein 7 (DNAH7). DNAH7 has a 13-residue insert between H2 and H3 compared to cytoplasmic dynein called the flap. Full MTBD sequence alignment in Figure S6A **B** – A model of DNAH7 (violet) was refined into the SRS^+^-DNAH7_2758-2896_ cryo-EM density (grey, lowpass filtered to 5Å). **C** – A comparison between the microtubule bound DNAH7 model (violet) and the low-affinity Flagellar dynein C NMR structure ((Kato et al., 2014), 2RR7, ensemble chain 8, docked to the same density, white). As with cytoplasmic dynein, the only major conformational changes between the two is the upward movement of CC1/H1 (purple and grey for DNAH7 and 2RR7 respectively) **D** – The SRS^+^-DNAH7_2758-2896_ cryo-EM density (MTBD colored purple) thresholded at a low level, revealing the extension of the flap to contact the adjacent protofilament **E** – Rotation of the DNAH7 MTBD relative to cytoplasmic dynein-1. The tubulin was aligned in the cytoplasmic and axonemal dynein models. This reveals a rotation around the base of H6/CC2 towards the adjacent protofilament in DNAH7 (violet) compared to cytoplasmic dynein-1 (white). **F** – Close up view of the flap of the DNAH7 MTBD model with its corresponding density. **G** – Corresponding view to F, without the electron density. **H** – The DNAH7 map lowpass filtered to 10Å reveals a new connecting density that cannot be explained by the flap (DNAH7 model docked, violet). **I** – The extra density observed in H is not seen in the cytoplasmic dynein-1 density filtered to the same resolution **J** – The C-terminal tail of β-tubulin could potentially explain the density in H, given its length and possible binding site in a positively charged pocket on the top surface of DNAH7 (surface colored by coloumbic potential) **K** – Cytoplasmic dynein-1 (surface colored by coloumbic potential) does not have the same positive patch on DNAH7, suggesting that the C-terminal tail would not dock in the same way.

A previous NMR study of *Chlamydomonas* flagellar dynein-c reported a low microtubule affinity (Kato et al., 2014). In contrast, another study observed that single dynein-c molecules bind well enough to move along microtubules (Sakakibara et al., 1999), and an artificial construct fusing the DNAH7 MTBD onto a cytoplasmic dynein-1 stalk bound to microtubules with high-affinity (Imai et al., 2015). We initially expressed and purified a 12-heptad stalk-SRS DNAH7 MTBD construct, but observed poor microtubule decoration in EM. We therefore made an SRS-fusion containing the mouse cytoplasmic dynein-1 stalk and the human DNAH7 MTBD. Consistent with (Imai et al., 2015) this construct (SRS^+^-DNAH7_2758-2896_) exhibited strong decoration of microtubules in cryo-electron micrographs (**Figure S1A**). We subjected these decorated microtubules to the same routine of data collection as before (**Table 1**).

Following 3D classification and refinement, we obtained a 4.8Å resolution map (**Figure 3B, S4A, S7A**). A model of the DNAH7 MTBD was refined into the density using a homology model to the low-affinity NMR structure (PDB 2RR7, (Kato et al., 2014)) as a starting point (**Figure 3B, S7B, Table 2**). Comparing the final microtubule-bound model to the original low-affinity structure, we observe the same conformational changes as for SRS-DYNC1H1_3260-3427_ (**Figure 3C**). Namely, the majority of the MTBD remains unchanged, but H1 and CC1 move up into a raised position over the intradimer interface. Furthermore, aligning our cytoplasmic dynein-1 model to the DNAH7 model shows that they adopt almost identical conformations (**Figure S7C**). Given that these are two distantly related dyneins, this suggests that the fundamental structural basis for microtubule binding is universal.

The biggest difference between the cytoplasmic dynein-1 and DNAH7 models lies in the flap. At lower threshold levels an elongated density emerges from the DNAH7 MTBD and contacts the adjacent protofilament (**Figure 3D + S7B**). There are two contacts between the flap and this protofilament, corresponding to H10 of β-tubulin and loop H6/7 of α-tubulin (**Figure 3F/G**). As such, DNAH7 contacts four tubulin subunits at once when it binds to the microtubule. On account of its appearance only at lower threshold levels, we conclude that the flap has a degree of flexibility.

Another difference between the two structures is the orientation of the MTBD on the microtubule. Aligning the tubulin of the DNAH7 and cytoplasmic dynein-1 models reveals that the DNAH7 MTBD is tilted relative to cytoplasmic dynein (**Figure 3E**). This can be described by a 7° rotation around the microtubule contact site at the base of H6. DNAH7 is tilted in the same direction as the flap, suggesting that its interaction with the adjacent protofilament pulls on the entire MTBD.

Lowpass filtering and thresholding the map to a lower level revealed the presence of an additional link between the DNAH7 MTDB and the adjacent protofilament (**Figure 3H**). Due to its proximity to the C-terminus of β-tubulin, we suspect that this density is attributed to the β-tubulin “E-hook”. This is a normally unstructured ~20 residue chain of mostly glutamate residues that strongly contributes to the electronegative surface of microtubules (Nogales et al., 1998; Redeker et al., 1992). We note that the same link is not observed in our cytoplasmic dynein-1 density (**Figure 3I**) at any threshold. Examination of the surface charges of our two models reveals a large positively charged patch on the top of the DNAH7 MTBD that is absent in cytoplasmic dynein-1 (**Figure 3J/K**). Modelling of the β-tubulin E-hook show that it is capable of reaching this patch. As such, we conclude that on top of a core microtubule interaction shared with cytoplasmic dynein, DNAH7 is bolstered by two links to the adjacent protofilament.

### DNAH7 binding to microtubules induces changes in microtubule cross section

During the process of solving the DNAH7 MTBD structure, we noticed that the microtubule was distorted compared to our other reconstructions. Initial application of local symmetry, using the same operators as our previous structures, resulted in a weaker decorating density and blurring of tubulin helices (**Figure S7D/E**). This suggested that the averaging between different protofilaments was incoherent, and the cross-section of the microtubule was distorted. We therefore used Relion to refine the local symmetry operators, which improved the features and led to our final 4.8Å map (**Figure S7F**).

To explore the distortion further, we manually measured the rotation between each protofilament relative to the long axis of the microtubule (**Figure 4A**). Measurements for the SRS-DYNC1H1_3260-3427_ map were as expected, with the angle between each protofilament close to 27.69° (360°/13 protofilaments, S.D.=0.17°) (**Figure 4B**). In contrast, our refined SRS^+^-DNAH7_2758-2896_ structure exhibited protofilament angles ranging from 26.5° to 28.7° (S.D=0.75°, **Figure 4B**), representing large changes in the local curvature of the microtubule.

**Figure 4.**
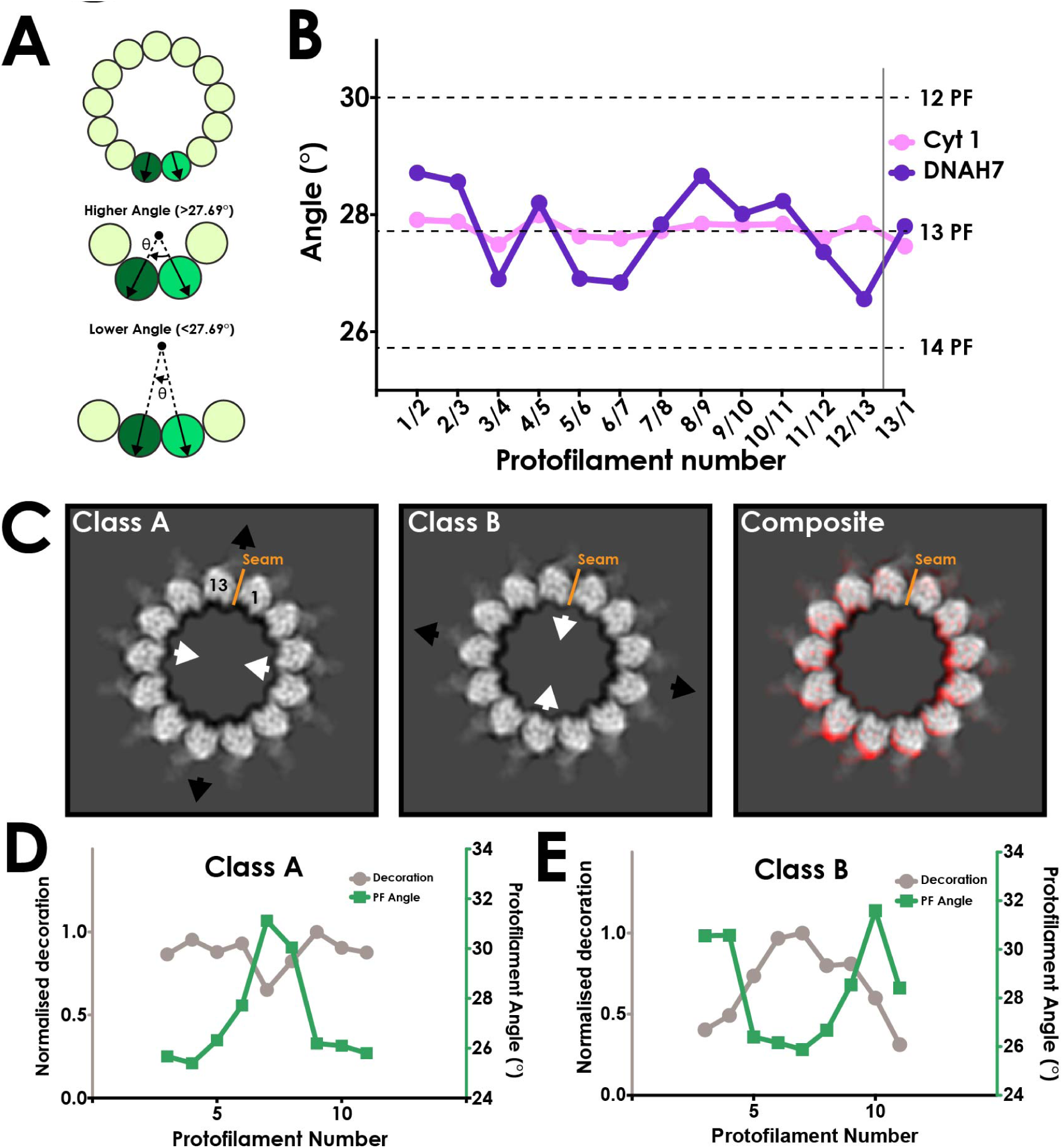
**A** – Schematic representing measurement of protofilament angles. A PDB model for tubulin (PDB 5SYF) was docked into adjacent protofilaments, and the rotation to superimpose the two models was measured. Local curvature between each neighbouring protofilament can thus be measured to match a canonical 13-protofilament microtubule (27.69° rotation), or be more or less curved. **B** – A plot of the angle between each neighboring pair of protofilaments in the cytoplasmic dynein-1 (Cyt 1, pink) and DNAH7 (purple) maps prior to symmetrization. In DNAH7, the microtubule has been distorted from an perfect circle. Protofilament angles from canonical 12-PF (30°), 13-PF (27.69°) and 14-PF (25.71°) are shown, and the position of the seam is indicated by the grey line **C** – A projection of one helical rise of Class A and Class B (left and center) with arrows indicating distortion in the curvature of the microtubule. Note that Class B and Class C are squashed and extended in reciprocal directions. (Right) A composite image showing the distortion between Class A (Red) and Class BC (white) **D** – A plot of the protofilament angles (green) and relative decoration (grey) on each protofilament for class A. **E** – A plot of the protofilament angles (green) and relative decoration (grey) on each protofilament for class B.

Our initial 3D classification of the SRS^+^-DNAH7_2758-2896_ particles resulted in a single class used in the refinement. However, to investigate the curvature further we reclassified the SRS^+^-DNAH7_2758-2896_ data with modified classification parameters (see methods, **Figure S7H**). This resulted in two extreme classes, “A” and “B”, with large differences in cross-sectional curvature (**Figure 4C, Movie 1**). The formation of multiple good classes indicates that the microtubules in this dataset form a continuous distribution of distorted curvatures. Class A and B had even greater local cross-sectional curvature distortions than our refined map, with protofilament angles ranging from 25.1° to 31.8° (**Figure 4D/E**). The local curvatures between protofilaments that result from these distortions are thus within the range of those seen in canonical 12 (25.71°) and 14 (30°) protofilament microtubules.

These distortions change the microtubule cross-section from a circular to an elliptical profile. The ellipticity (the ratio between the long and short diameters of an ellipse) of class A and B is 0.936 and 0.942 respectively (**Figure S7I/J**). This is much greater than microtubule ellipticity measured both in our cytoplasmic dynein-1 structure (**Figure S7K**) and previous studies (Kellogg et al., 2017), indicating that DNAH7 is responsible for this distortion.

To determine how DNAH7 binding affects the curvature of the microtubule we measured the relationship between the local curvature of the microtubule and the level of MTBD decoration in classes A and B. We observe a clear correlation in which the decoration is highest at low local protofilament curvatures (**Figure 4D/E, S8**). Based on this localization, we propose that the DNAH7 MTBD induces a local flattening of the microtubule. The pattern of decoration also suggests that the DNAH7 MTBD binds more weakly to higher curvatures and more strongly to flatter curvatures.

Given that the only major difference between the cytoplasmic dynein-1 and DNAH7 MTBD is the flap, it seems likely that this is the element responsible for the flattening. This is supported by our observation that the flap contacts the adjacent protofilament to the MTBDs binding site. Furthermore, the movement of the MTBD towards the adjacent protofilament suggests that there is tension created by the flap. We propose that this tension results in the relative movement between protofilaments that generates a local flattening.

## Discussion

### Using Relion to solve high-resolution structures of decorated pseudosymmetric microtubules

The workflow we present for solving microtubule structures in Relion is straightforward and does not require expert knowledge. To test the general applicability for decorated microtubules, we re-processed EMPIAR dataset 10030, which comprises EB3-decorated microtubules. This data set was previously refined to 3.5Å resolution using an iterative seam-finding protocol (Zhang and Nogales, 2015; Zhang et al., 2015). Following the Relion pipeline with the same data resulted in an asymmetric map with lower levels of density on the protofilaments either side of the seam, suggesting that not all particles were aligned correctly with respect to the seam (**Figure S3D/E**). However, after symmetrization the map extends to 3.5Å resolution, and is essentially identical in appearance to the published structure (**Figure S3A-C**). We conclude that Relion can be used to reconstruct pseudosymmetric microtubules and their decorating partners to high resolution.

We note that many of the previously introduced methods for microtubule reconstruction explicitly determine, and enforce, a single seam orientation for each microtubule (Sindelar and Downing, 2007; Zhang and Nogales, 2015). In Relion, the in-plane rotation, translations, and out-of-plane rocking (tilt) angle can be restrained to similar values in neighbouring segments from each microtubule through the use of Gaussian priors. However, such a prior has not been implemented for the first Euler angle, i.e. the rotation around the helical axis. Therefore, each segment is aligned independently from its neighbours in the same microtubule, and mistakes in alignment of the seam can be made on a per-segment basis. Therefore, at least in its current implementation, Relion appears to require a larger decorating density (such as the dynein MTBD) to fully align the seam.

### A universal high-affinity state of the dynein MTBD

Our new 4.1Å reconstruction of the cytoplasmic dynein-1 MTBD allows us to identify the structural transitions that occur upon dynein binding to microtubules. The MTBD interface made up of H2, H3, H4 and H6 shows only minor changes upon microtubule binding, suggesting it is in a binding-primed conformation regardless of nucleotide state. This interface can initiate contact with the microtubule, but a stable microtubule-bound state cannot be achieved when dynein is in a low-affinity state due to a steric clash between H1 and β-tubulin. When the motor switches to a high microtubule affinity nucleotide state, H1 moves up and the MTBD can be further stabilized on the microtubule by interactions between H1 and β-tubulin. Conversely, when ATP binds to the motor, changes in the stalk (Schmidt et al., 2015) push H1 down and release the MTBD from the microtubule.

Combining our current work with previous studies (Carter et al., 2008; Kato et al., 2014), we now have structures of two distantly related dynein MTBDs both on and off the microtubule. In both the cytoplasmic dynein-1 and DNAH7 MTBDs the predominant change is the position of H1, suggesting this is a universal mechanism for controlling dynein binding to microtubules. However, the addition of an extended flap between H2 and H3 in DNAH7 results in important differences to cytoplasmic dynein. The flap extends the contact area with the microtubule to include 4 tubulin subunits, supplementing the core interaction observed in cytoplasmic dynein-1. We propose that this interaction drives large distortions in lattice curvature. We assign a second link to the adjacent protofilament to the acidic C-terminal tail of beta-tubulin binding to a pocket of positively charged residues on the top surface of the DNAH7 MTBD.

### DNAH7 induced distortions of the microtubule

Some members of the kinesin family of microtubule motors are known to induce changes in the microtubule lattice. These include a longitudinal extension by kinesin 1 (Peet et al., 2018) and the peeling back of single protofilaments by kinesin 13 (Hunter et al., 2003). Both these effects are related to the longitudinal axis however, and no effects on cross-sectional curvature have been observed to date. In contrast, some microtubule-associated proteins are sensitive to the lateral curvature of the microtubule. Doublecortin (DCX) and EB3 both bind in the cleft between two adjacent protofilaments and contact four tubulin subunits (Moores et al., 2004; Zhang et al., 2015). These lateral interactions promote formation of 13-protofilament microtubules during polymerization. This promotes a rounder, more regular lattice. Therefore, DNAH7 is unusual in directly distorting the cross-sectional curvature of mature microtubules.

A number of other axonemal dyneins possess an MTBD flap (**Figure S6B/C**). In *C. reinhardtii* the γ outer arm dynein and the a, b, c, d and e inner arm dyneins all possess an 18/19-residue loop between H2 and H3 (**Figure S6C**). Cryo-electron tomography has mapped the position of each dynein MTBD in the axoneme (**Figure 5A**) (Bui et al., 2008; Liu et al., 2008; Nicastro et al., 2006; Song et al., 2018). Microtubules in the axoneme form a doublet structure, consisting of a 13-protofilament A-tubule and a connected 10-protofilament B-tubule. The tail of axonemal dyneins dock onto the A-tubule, positioning the MTBDs to bind to an adjacent doublet B-tubule. Strikingly, the flap containing dyneins are spread along the length of the axonemal repeat but only contact two pairs of protofilaments on one side of the B-tubule (**Figure 5B**). As such, there is one side of the B-tubule on which flap-containing dyneins act at a high local concentration. This is in contrast to our DNAH7 decorated microtubule structure, in which near-saturation binding around the entire microtubule may confound local effects of individual DNAH7 MTBD binding. The effect of dynein binding to microtubules in the axoneme may therefore result in even greater distortions than those we observed.

**Figure 5.**
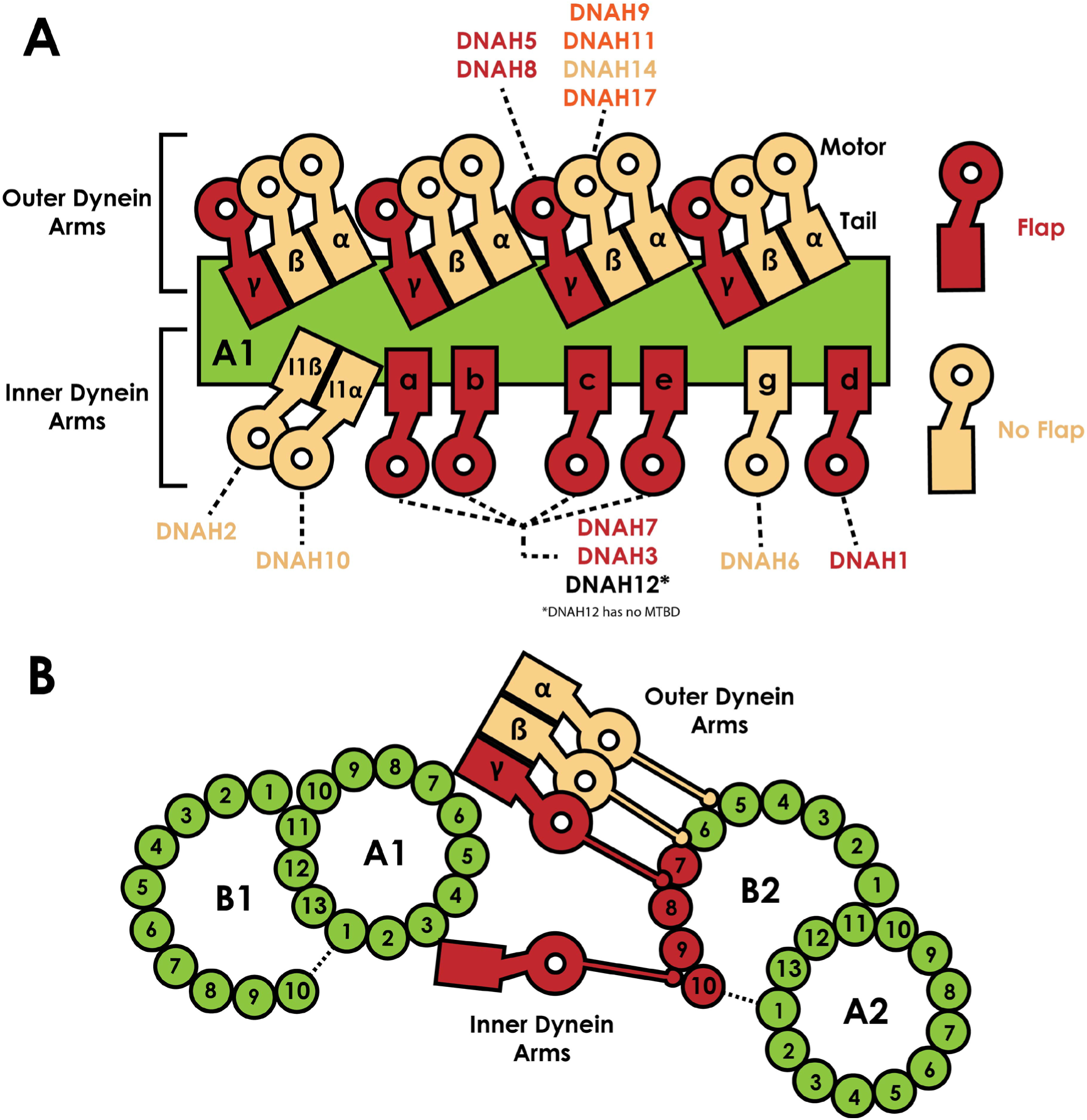
**A** – Schematic view of the positions the dynein heavy chain tails dock on the A-tubule of the microtubule doublet (green) in the 96nm axonemal repeat, based on previous structural data on C.reinhardtii and Sea urchin sperm flagella (Bui et al., 2008; Lin and Nicastro, 2018; Nicastro et al., 2006). Heavy chains are named according to the C. reinhardtii nomenclature, and the human orthologues are linked. Heavy chains are color coded based on whether they possess a MTBD flap (Red) or not (beige). **B** – Orthogonal view of A, now looking down the flagella. Links between the inner and outer dynein arms and the exact protofilament of the adjacent doublet has also been established by structural work (Song et al., 2018). The position of the tails on tubule A1 positions the MTBDs on tubule B2. We note each of protofilaments 7 to 10 on tubule B2 are contacted by flap-containing dynein MTBDs.

Axonemal dyneins drive ciliary bending (King, 2018; Satir et al., 2014). Most models suggest that localized activation and inhibition of axonemal dyneins is needed to create the imbalance in forces across the axoneme that result in an overall bend. These models rely on communication between axonemal dyneins such that they are active or inactive at the right time. We speculate that the flap could play a role in communication. We observed higher decoration at lower lateral curvature in our classes (**Figure 4D/E**), suggesting that the DNAH7 MTBD could act as a curvature sensor in the axoneme. The cross-section of microtubules is thought to flatten during bending (Memet et al., 2018), in which case dynein binding would cyclically change during the axonemal beat. Alternatively, the flap-induced distortion of the microtubule could induce cooperative binding of adjacent dyneins, potentially helping a waveform spread through the cilium. Higher-resolution structures of axonemal dyneins in the context of a ciliary beat will be required required to test these hypotheses.

## Acknowledgements

This study was supported by the MRC-LMB EM facility. We thank C. Sindelar for initial help with microtubule processing; C. Savva, G. Cannone, J. Grimmett, and T. Darling for help and technical support at the MRC-LMB; S. Neumann and J. Gilchrist for technical support at Diamond; F. Abid Ali and C. Lau for comments on the manuscript. We acknowledge Diamond for access and support of the Cryo-EM facilities at the UK national electron bio-imaging centre (eBIC), (proposal EM17434), funded by the Wellcome Trust, MRC and BBSRC. This work was funded by grants from the Wellcome Trust (WT210711) and the Medical Research Council, UK (MC_UP_A025_1011) to A.P.C.

## Author Contributions

S.L. prepared samples, collected data and performed all cryo-EM processing. S.H. and S.H.W.S implemented non-crystallographic symmetry in Relion and provided advice. A.P.C supervised the project. S.L. and A.P.C wrote the manuscript with input from S.H.W.S.

## Data deposition

Cryo-EM maps have been deposited in the Electron Microscopy Data Bank (EMDB) under accession numbers EMDxxxx. Refined atomic models have been deposited in the Protein Data Bank (PDB) under accession numbers xxxx.

## Materials and Methods

### Protein Preparations

6xHis 12-heptad SRS fusion proteins were expressed in SoluBL21 *Escherichia coli* cells (Invitrogen) from a pet42a vector. The mouse SRS-DYNC1H1_3260-3427_ construct is identical to the SRS-MTBD-85:82 used in (Carter et al. 2008). The DNAH7 MTBD and stalk was made as a synthetic gene product (EpochGene). SRS^+^-DNAH7_2758-2896_ was made by cloning the DNAH7 sequence into SRS-DYNC1H1_3260-3427_ in the place of the MTBD, delineated by the universally conserved proline residues as in (Imai et al. 2015). Cells were grown in LB media at 37°C until their OD_600_ measured 0.4-0.6, at which point they were supplemented with 1mM IPTG and grown for 16 hours at 16°C. Cultures were spun at 4000x rcf for 15 minutes, and used directly for purification. Both SRS constructs were purified according to the same protocol.

A 1L pellet was resuspended in 50mL Lysis buffer (50mM Tris pH8.0, 100mM NaCl, 1mM MgCl_2_, 10% Glycerol, 10mM Imidazole pH8.0, 1mM DTT, 2mM PMSF) and lysed by sonication. The lysate was centrifuged at 30000x rcf in a Ti70 rotor (Beckman) for 30 minutes and at 4°C. The supernatant was loaded onto a 5mL NiNTA HisTrap HP Column (GE), washed with 10 column volumes of 10% elution buffer (Lysis buffer with 500mM Imidazole pH 8.0 and without PMSF) and eluted with a step gradient to 40% elution buffer. Peak fractions were pooled and concentrated in a 15mL 30kMWCO centrifugal concentrator (Amicon) to a concentration of ~5mg/mL. Aliquots were snap frozen in liquid nitrogen.

ZZ-tagged human cytoplasmic dynein 1 motor domain (DYNC1H1_1230-4646_) was cloned into pFastBac and expressed in *Sf9* insect cells as in (Schmidt et al. 2015). A 1L pellet was resuspended in 50mL ZZ-Lysis buffer (as above but without imidazole) and dounce homogenised with 30 strokes. Lysate was centrifuged at 500000x rcf in a Ti70 rotor (Beckman) for 60 minutes and at 4°C. Supernatant was mixed with 2mL IgG Sepharose 6 Fast Flow resin (GE, equilibrated in ZZ-lysis buffer) on a horizontal roller for 2 hours at 4°C. The mixture was applied to a gravity flow column, and the resin was washed with 150mL ZZ-Lysis buffer and 150mL TEV buffer (50mM Tris pH 7.4, 150mM KOAc, 2mM MgAc, 1mM EGTA, 10% Glycerol, 1mM DTT). The resin was resuspended in 5mL TEV buffer, supplemented with 0.1mg/mL TEV protease and incubated on a horizontal roller at 25°C for 80 minutes. The sample was reapplied to a gravity flow column, the eluate was collected and concentrated to 6mg/mL with a 15mL 100kMWCO centrifugal concentrator (Amicon) and snap frozen in aliquots.

Aliquots of each sample were gel filtered prior to each grid freezing session. Thawed sample was spun through a 0.22um spin filter (Amicon) to remove aggregates and loaded onto a Superose 6 10/300 gel filtration column (GE) equilibrated in GF buffer (25mM Tris pH8.0, 50mM NaCl, 1mM MgCl_2_, 1mM DTT). Peak fractions were pooled and concentrated in a 4mL 30MWCO Amicon centrifugal concentrator to 1/10^th^ of the original volume. The sample was then diluted 5-fold in salt-free GF buffer (i.e. without 50mM NaCl) and reconcentrated. This was repeated twice, resulting in a 25-fold dilution of the NaCl. The sample was further diluted to a final concentration of 2mg/mL to be used for grid freezing.

Lyophilised tubulin was resuspended in MES-NaCl buffer (25mM MES pH6.5, 70mM NaCl, 1mM MgCl_2_, 1mM DTT) to a concentration of 10mg/mL and snap frozen in aliquots. For polymerisation, an aliquot was thawed and mixed 1:1 with MES-NaCl buffer supplemented with 6mM GTP, and incubated at 37°C for 2 hours. 100uL MES-NaCl buffer supplemented with 20uM Taxol and pre-warmed to 37°C was added, and the sample was left at room temperature overnight. Before use, the microtubules were spun at 20000x rcf for 10 minutes, and resuspended in MES-NaCl buffer with taxol.

### Grid preparation

Quantifoil R1.2/1.3 Au300 grids were glow-discharged for 40 seconds. 4uL 0.4mg/mL microtubules was added to the grid and incubated at room temperature for 1 minute. This was removed by side blotting, 4uL of dynein was added and the grid was incubated for a further 2 minutes. Manual side blotting was repeated, and after the second MTBD application the grid was taken into the humidity chamber of a Vitrobot Mark II set to 100% humidity and 22°C. After 2 minutes, the grid was double-side blotted for 4 seconds and plunged into liquid ethane.

### Cryo-electron microscopy

Cytoplasmic dynein 1 MTBD-SRS grids were imaged on our in-house Titan Krios microscope, and DNAH7 MTBD grids were imaged on Krios III at Diamond eBIC. For cytoplasmic dynein, 1995 1.5s exposures were collected with a pixel size of 1.04Å^2^ and a fluence of 40e^−^/Å^2^s on a Falcon III detector in linear mode. For DNAH7, 4641 1.5s exposures were collected with a pixel size of 1.085Å^2^ and a fluence of 45e^−^/Å^2^s. Dynein motor domain decorated microtubules were imaged on a Polara microscope, with 2455 1.5s exposures collected with a pixel size of 1.34Å^2^ and a fluence of 37e^−^/Å^2^s on a Falcon III detector in linear mode. In each case, images were acquired with a defocus ranging between −1.5μm and −4.5μm semi-automatically in EPU.

### Image Processing

All processing was performed inside the Relion 3.0 pipeline (Zivanov et al. 2018). Details are given for processing the SRS-DYNC1H1_3260-3427_ data, followed by modifications to this workflow used for the other datasets. The unaligned raw movies were aligned and dose weighted in Relion’s implementation of MotionCorr2 using 4×4 patches (Zheng et al. 2017). CTF determination was performed with Gctf on dose-weighted micrographs (K. Zhang 2016). Manual picking and 2D classification was performed to generate references for autopicking. Start and end coordinates of 30 microtubules from 5 micrographs were extracted into 82Å segments (box size 512), resulting in ~650 particles which were classified into 5 classes. These were used as references for autopicking on all the micrographs, using the following parameters: mask diameter 497Å, in-plane angular sampling 1°, lowpass references 20Å, picking threshold 0.04, minimum inter-particle distance 79Å, maximum stddev noise 1.4, shrink factor 0.5, helical picking, tube diameter 400Å, helical rise 82Å, number of asymmetrical units 1, maximum curvature 0.4. Particles were extracted (4x binned) and entered for 2D classification into 100 classes. Classes were rejected if they were obviously not microtubules (carbon, ice etc), if they appeared blurred or poorly aligned, if they had low levels of decoration and if they showed signs of non-13-PF architectures (**Figure S1C)**. A 3D reference was made by docking a model of dynein MTBD decorated tubulin into density for a 13-PF microtubule (PDB 3J1T and EMD 6351 respectively). The PDB was converted to electron density in EMAN2 (*pdb2mrc)*. 3D classification of unbinned particles into 3 classes was used to separate out the remaining sample heterogeneity. The single good class was entered into a 3D refinement using the following parameters: initial angular sampling of 0.9°, and initial offset range and step sizes of 3 and 1 pixels respectively. C1 symmetry, inner tube diameter 100Å, outer tube diameter 400Å, angular search range tilt 15°, psi 10°, tilt prior fixed, range factor of local averaging 4, helical symmetry with 1 asymmetric unit, initial rise 82Å, initial twist 0°, central Z length 40%, local searches of symmetry, rise search 78-86Å, step size 1Å, no search for twist.

A solvent mask and 13-fold local symmetry was applied during refinement. For local symmetry, a mask was made by docking copies of PDB 3J1T into the protofilament to the left of the seam (if the MT is being viewed plus-end up). The PDB protofilament was then converted to electron density with the EMAN program *pdb2mrc*. This was converted into a mask with *relion_mask_create*. *relion_local_symmetry* requires a STAR file containing the translational and rotational operators needed to move the original mask onto each successive protofilament. The psi angle, rotating around the microtubule long axis, is given as multiples of −27.69° (360°/13). The center of rotation is the center of the microtubule lumen, so the only translation needed is the rise between adjacent protofilaments. For a 3-start helix, there is a rise of 1.5 dimers through 360°. The refined helical rise between dimers in the same protofilament as measured by Relion was 82.29Å. As such, the _rlnOriginZ parameter increases by multiples of 9.495Å (82.293 * 1.5 / 13). Local symmetry was applied during refinement with the additional argument --local_symmetry. Following completion of refinement, local symmetry was applied to both unfiltered half maps. Postprocessing and resolution assessment was performed with three tubulin dimers docked along a single protofilament as previously (R. Zhang et al. 2015a; R. Zhang, LaFrance, and Nogales 2018).

Refinement of the dynein motor domain also followed this protocol. For the DNAH7 structure, initial 3D classification did not result in a coherent class. Instead all good particles following 2D classification were entered into 3D refinement, resulting in a map with blurred features. Following this, 3D classification into 8 classes using the orientations used in the refinement (i.e. with no image alignment) was performed. Local symmetry operators were found for the resulting map with the search command in *relion_localsym* (see https://www2.mrc-lmb.cam.ac.uk/relion/index.php?title=Local_symmetry). Classes A and B were made in another classification job, this time with Tau=20 and using 0.9° local angular searches.

The EB3 dataset was downloaded from EMPIAR (ID 10030) and processed as for the dynein MTBD with modifications. 3D classification was skipped since the microtubules in this dataset almost exclusively have 13 protofilaments (R. Zhang et al. 2015b). EB3 does not bind across the seam, which means that applying regular 13-fold symmetry was not appropriate. A separate mask was created for the tubulin and EB3 densities. The tubulin mask and EB3 masks were applied with 13- and 12-fold symmetry respectively.

Local resolution estimation was performed in *relion_postprocess*.

### Model Building

For cytoplasmic dynein 1 a low-affinity crystal structure (PDB 3ERR) was used as the starting model for refinement. For human DNAH7, a sequence alignment to *C. reinhardtii* flagellar dynein c was used to generate a homology model to the low-affinity NMR structure (PDB 2RR7) in Modeller (Sali and Blundell 1993). The homology model was used as an initial model. The models were fit in their respective maps using Chimera (Pettersen et al. 2004). A tubulin dimer was also docked in to the density using PDB 5SYF (for SRS^+^-DNAH7_2758-2896_ the tubulin dimer being contacted by the flap was added as well). Coot Real Space refine zone (Emsley and Cowtan 2004) was used to manually fit the model to the density, followed by whole model refinement using Phenix real space refine (Adams et al. 2010). These two steps were performed iteratively until the model to map measures were maximised. For model to map FSC curves (**Figure S4B**), phenix.mtriage was used. All model visualisation was performed in Chimera (Pettersen et al. 2004).

### Analysis of microtubule distortion

Protofilament angles were measured by docking a tubulin dimer pdb model (5SYF) into two adjacent protofilaments of the relevant reconstruction in Chimera. The relative rotation was measured with the “measure rotation” command. Ellipticity was measured with the Matlab fit_ellipse script deposited in the Mathworks file exchange (https://www.mathworks.com/matlabcentral/fileexchange/3215-fit_ellipse). x,y coordinates for each protofilament were obtained from maximum intensity projections of each class, binarised with the same threshold in FIJI (Schindelin et al. 2012). The center of mass of each protofilament was used as the coordinate.

---

**Figure S1.**
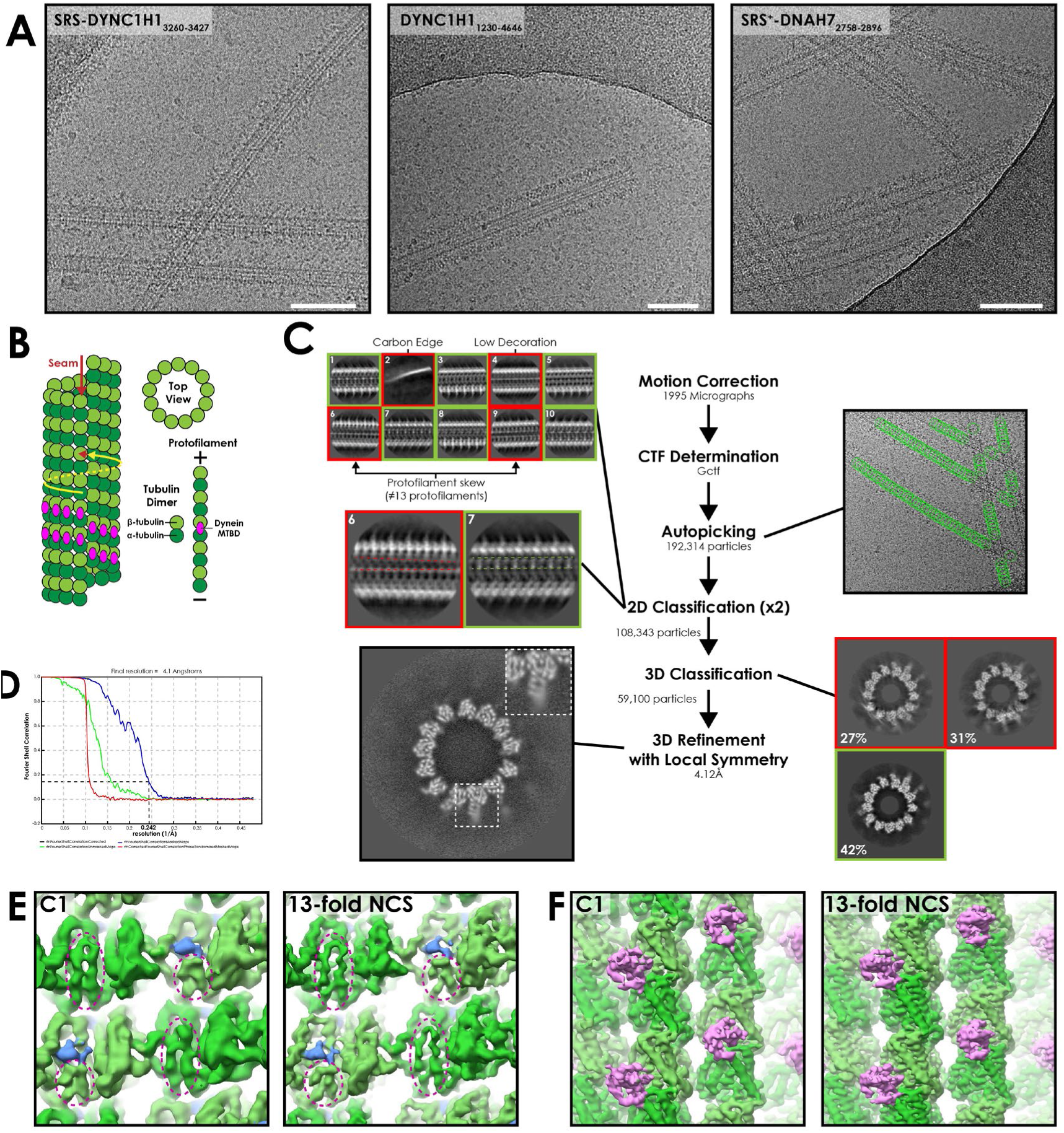
**A** – Representative cryo-EM images from each of the three datasets presented in this work. White bar corresponds to 100nm **B** – Cartoon depicting a 13-protofilament microtubule with a seam. α-tubulin (dark green) an β-tubulin (light green) form a constitutive dimer that polymerises end-on-end as a protofilament. 13-protofilaments are arranged in a ring to create a tube. Lateral interactions are like-for-like (α:α and β:β, yellow arrow) except at the seam (red arrows). This asymmetry means conventional helical averaging cannot be performed. The dynein MTBD (magenta) binds to the intradimer interface. **C** – Processing workflow for the cytoplasmic dynein-1 MTBD structure. Motion correction and CTF determination were performed as standard. Autopicking was optimised to pick all microtubules. 2D and 3D classification was used to remove bad particles. In 2D classification, bad classes included those with incorrect features (e.g. the carbon edge, class 2), low decoration on the side of the microtubule (class 4) or identifiable non-13 protofilament architecture (classes 6 and 9). The latter can be distinguished on account of the unique lack of protofilament twist in 13-protofilament microtubules (that is, protofilaments run exactly parallel to the microtubule long axis). This means that the repeating patterns in the classes should run parallel (class 7, expanded), whereas particles with protofilament twist will have converging or diverging patterns along the microtubule (class 6, expanded). This method is then supplemented with 3D classification to remove microtubule architectures with small protofilament skews that may not show up in short 2D segments, as well as particles with poor contrast or high-resolution detail. Refinement of the good class from 3D classification followed by 3D refinement with local symmetry resulted in a 4.1Å map (FSC_0.143_ cut-off, gold standard). **D** – Fourier shell correlation (FSC) curve for the refined SRS-DYNC1H1_3260-3427_ structure. **E** – Luminal view of the protofilament either side of the seam in the final C1 (asymmetric) and final symmetrized SRS-DYNC1H1_3260-3427_ maps. α- and β-tubulin (dark and light green) can be differentiated by the length of the S9-S10 loop (dotted circles). In α-tubulin the loop is extended, whereas in β-tubulin it is short, making room for the taxol binding site (taxol in blue). The asymmetry at the seam is clearly defined in both the C1 and symmetrized map **F** – Exterior view of the protofilament either side of the seam in the final C1 (asymmetric) and symmetrized SRS-DYNC1H1_3260-3427_ maps. MTBD density (magenta) undergoes an additional rise across the seam, reconfirming the definition of the seam.

**Figure S2.**
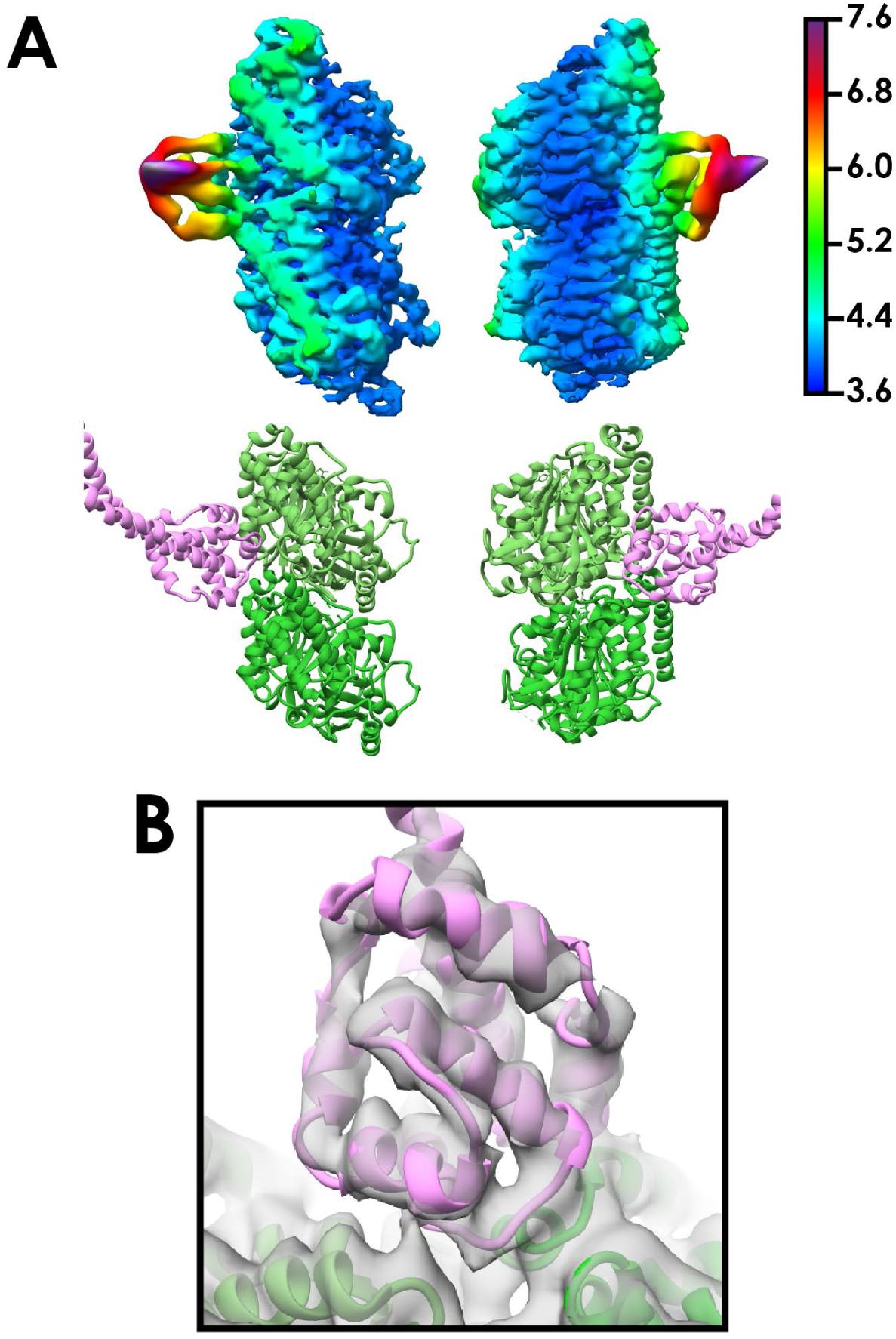
**A** – Density corresponding to one tubulin dimer and the cytoplasmic dynein-1 MTBD colored and filtered according to local resolution as determined in relion_postprocess. Density (top) and the corresponding views in the model (bottom) are shown. **B** – Orthogonal view of Figure 1D, showing the refined cytoplasmic dynein-1 MTBD docked into density lowpass filtered to 5Å.

**Figure S3.**
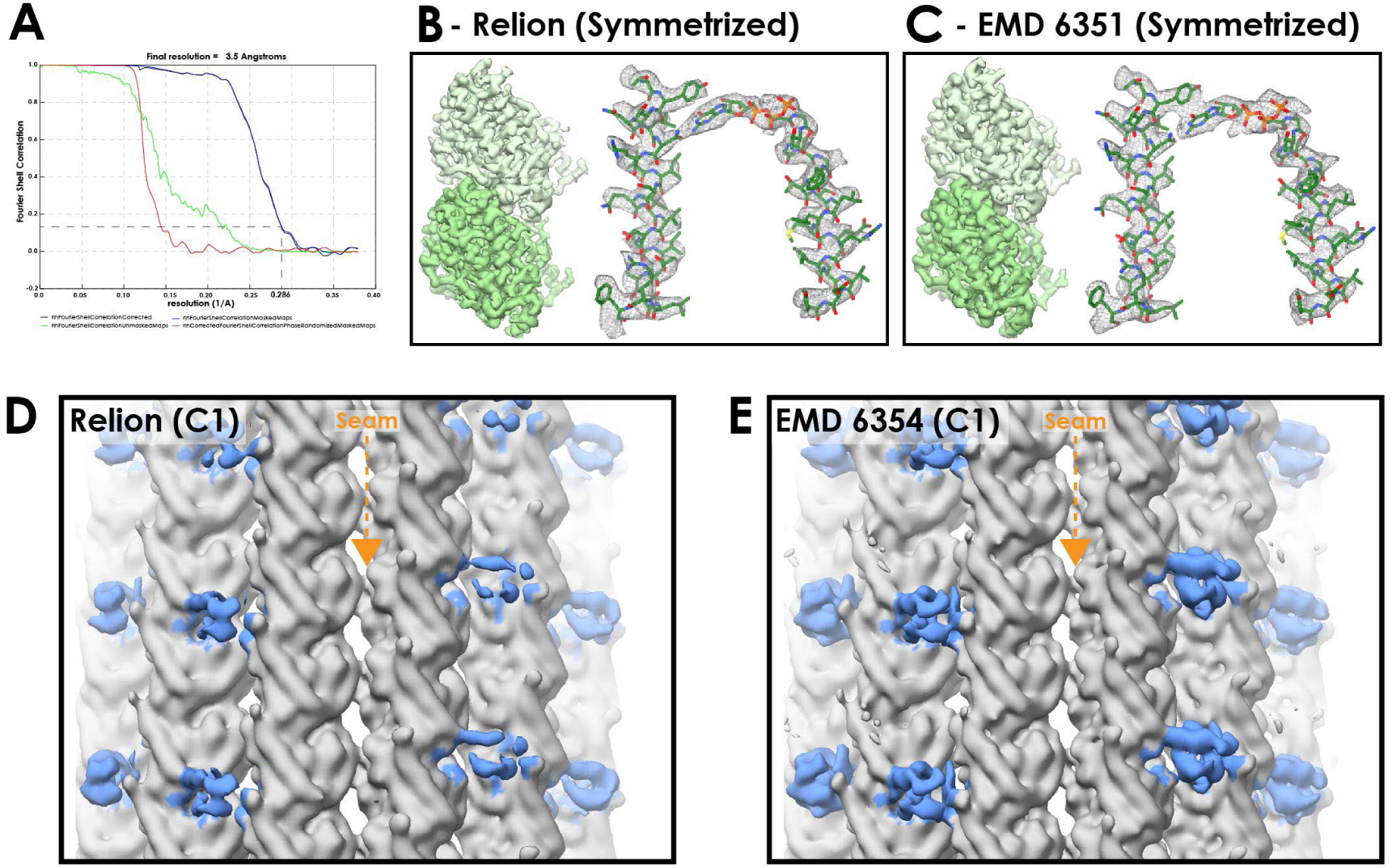
A – FSC curves for EMPIAR dataset 10030 (EB3 decorated microtubules) processed in the Relion pipeline following symmetrization (FSC_0.143_ cut-off, gold-standard) B – Density from the Relion refined EB3-microtubule dataset, following symmetrization. Mesh representation of alpha-tubulin H4+H7 and nucleotide. C – Corresponding views of the published map processed from the same dataset but refined with the seam-finding protocol (EMD 6351 – (Zhang and Nogales, 2015)). D – Density at the seam of the C1 (asymmetric) reconstruction from Relion (EB3 density colored blue). The EB3 density close to the seam is weak, suggesting that the seam is poorly aligned in some particles E – Corresponding view of a C1 reconstruction using the seam-finding protocol refinement (EMD 6354). The EB3 density is as strong around the seam as in neighbouring protofilaments.

**Figure S4.**
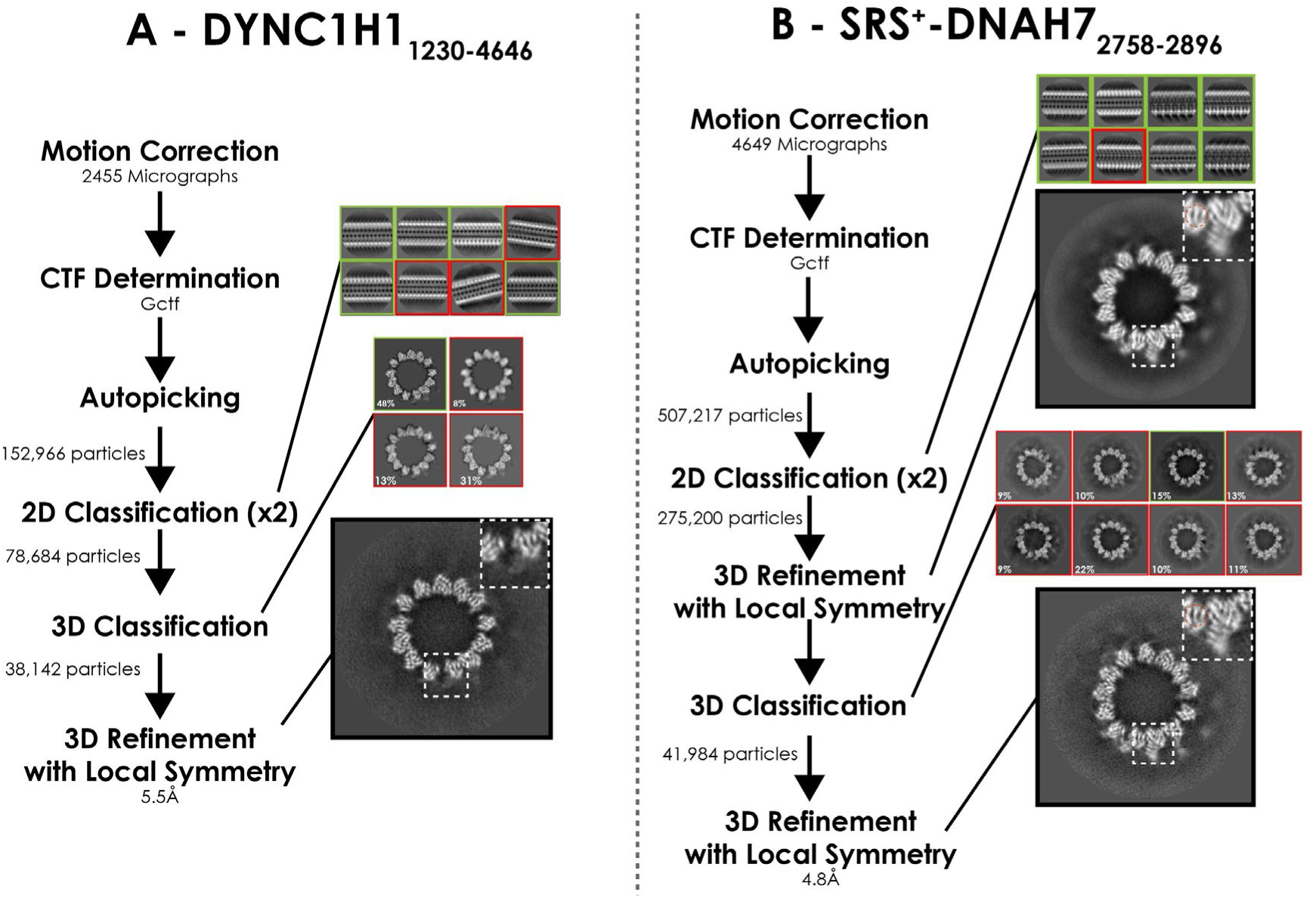
**A** – Processing pipeline for DYNC1H1_1230-4646_ structure **B** – Processing pipeline for DNAH7-MD structure. 3D classification directly after 2D classification did not result in a good class. 3D refinement was performed, and these orientations were used for 3D classification. This good class was taken forwards for 3D refinement, creating the final DNAH7 structure.

**Figure S5.**
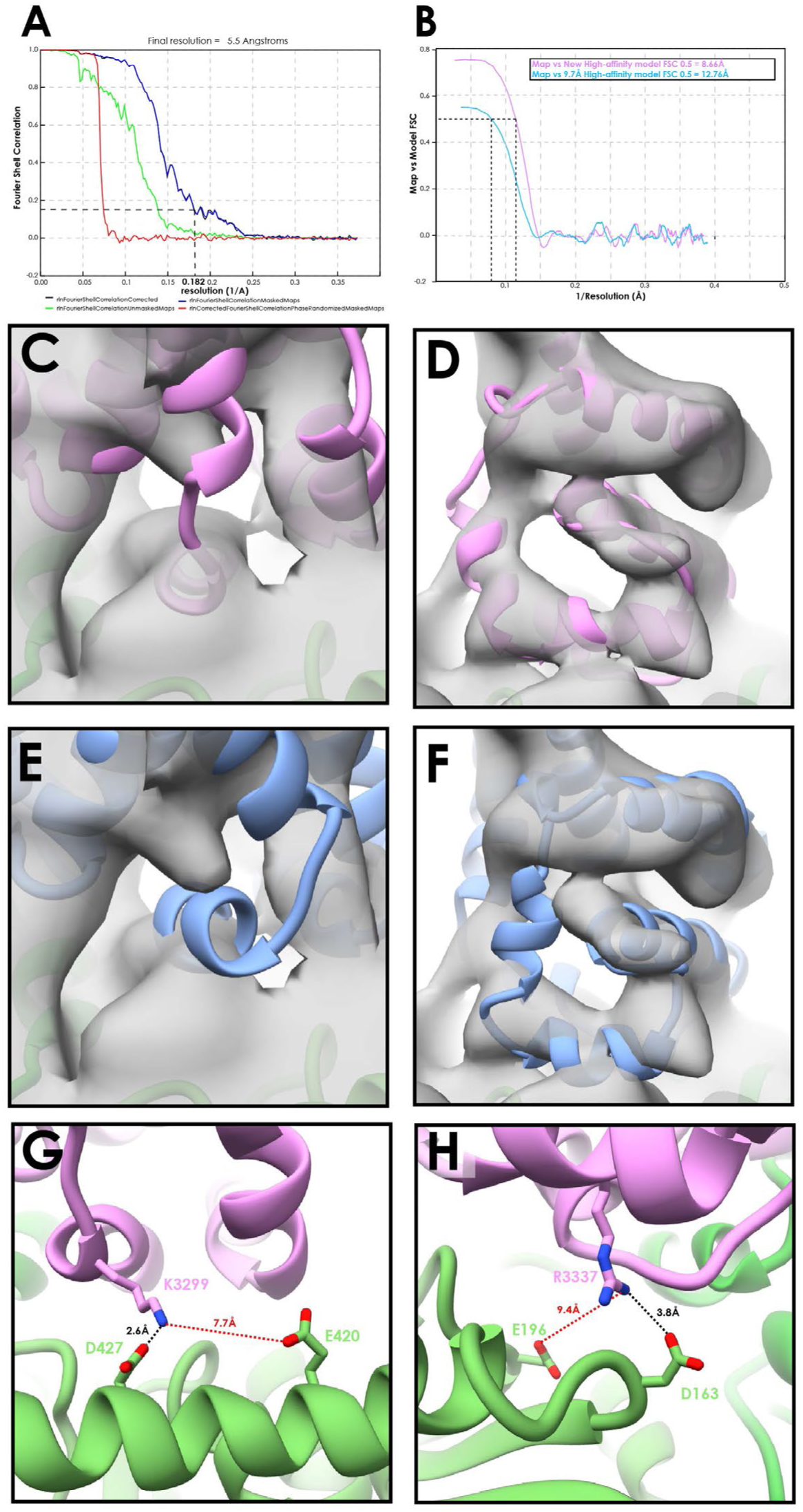
**A** – FSC curve for the DYNC1H1_1230-4646_ decorated microtubule structure (FSC_0.143_ cut-off, gold-standard) **B** – Model to map FSC curves for our new high-affinity cytoplasmic dynein-1 model or the previous 9.7Å high-affinity model (3J1T) docked into the DYNC1H1_1230-4646_ map **C** – Our newly refined cytoplasmic dynein-1 high-affinity model (Figure 1D/E) docked into the DYNC1H1_1230-4646_ map (filtered to 8Å), viewing CC1 and H1 **D** – Orthogonal view of C, viewing H2-H4 **E** – Equivalent to C, but with the previous 9.7Å cryo-EM MTBD model docked. H1 is now fully outside the density, compared to a good fit in B **F** – Orthogonal view of E, showing most of H2, H3 and H4 outside the density, compared to good fits in C **G** – The 4.1Å resolution density does not show side-chain positions for most of the MTBD. However, based on our model and all possible rotamer conformations a number of interactions between the MTBD and the microtubule identified previously (Redwine et al., 2012) are too far apart in the new model. The tubulin residue interacting K3299 was suggested to be E420, but this is now too far to interact, and we suggest that D427 is the more likely partner. **H** – Similarly, R3337 is now more likely to interact with D163 than E196

**Figure S6.**
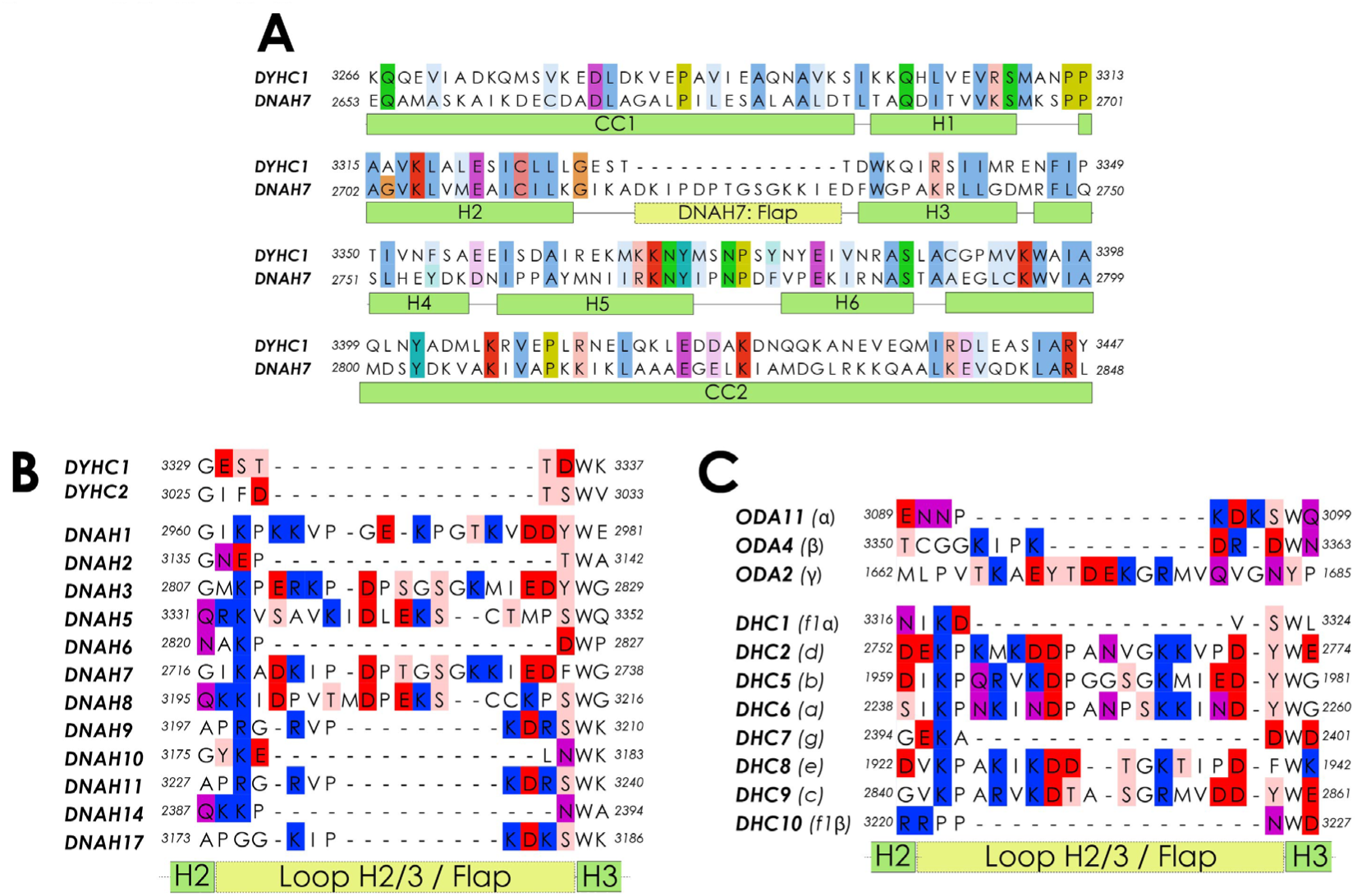
**A** – Full sequence alignment for the MTBD of human cytoplasmic dynein-1 (DYHC1) and human DNAH7 **B** – A sequence alignment of the region between H2 and H3 of the MTBDs of the human dynein heavy chains. DNAH1, 3, 5, 7 and 8 all have a 18/19 residue long flap, while all other dyneins have a much shorter loop. **C** – Equivalent to B for Chlamydomonas reinhardtii axonemal dynein heavy chain sequences.

**Figure S7.**
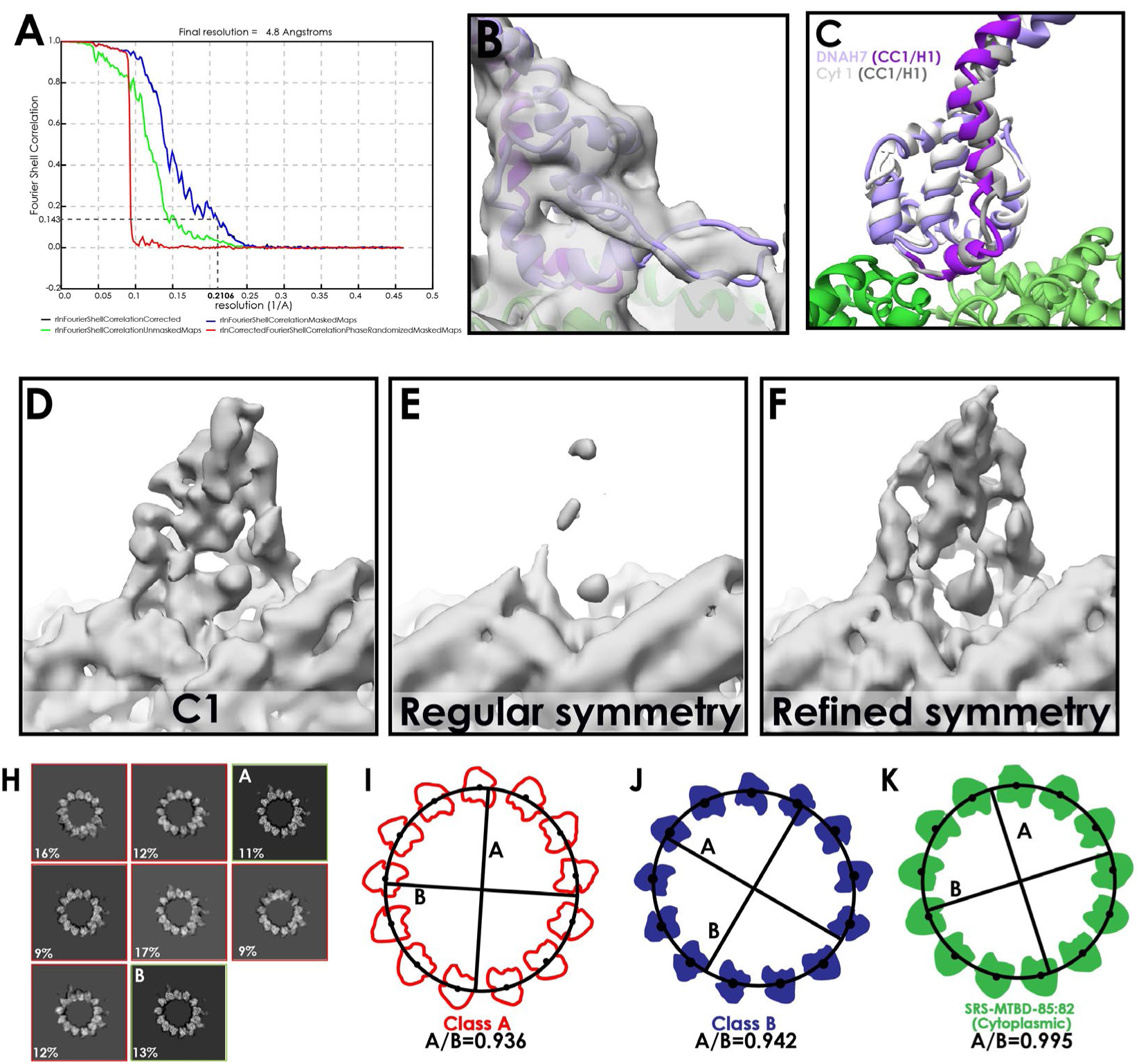
**A** – FSC curve for the final, symmetrized (with refined operators) DNAH7 map (FSC_0.143_ cut-off, gold-standard) **B** – The refined DNAH7 model docked into density, at a low threshold to visualize the flap **C** – A comparison of our newly refined high-affinity cytoplasmic dynein-1 and DNAH7 MTBD models. **D** – MTBD density from one protofilament of the asymmetric DNAH7 final map. **E** – Corresponding view to D, following symmetrization of the DNAH7 map using the regular, circular operators used for the previous structures. The decorating density is much weaker, suggesting that the averaging was incoherent **F** – Corresponding view to D, following symmetrization of the DNAH7 map using local symmetry operators refined in Relion. This finds the optimal position for each mask, and as such accounts for any deviations in curvature. The density is now much strong, and contains higher quality MTBD features than the asymmetric map. **G** – Binarised representation of classes B and C, aligned at a seam-adjacent protofilament. **H** – Results from 3D classification of DNAH7 when larger Tau value and local refinements were used (see methods), generating classes “A” and “B” (indicated) **I** – Class A with protofilament coordinates. An ellipse was fit to the coordinates (black line), from which the ellipticity is shown. **J** – As for H, but with Class B **K** – As for H, but with the refined SRS-DYNC1H1_3260-3427_ map

**Figure S8.**
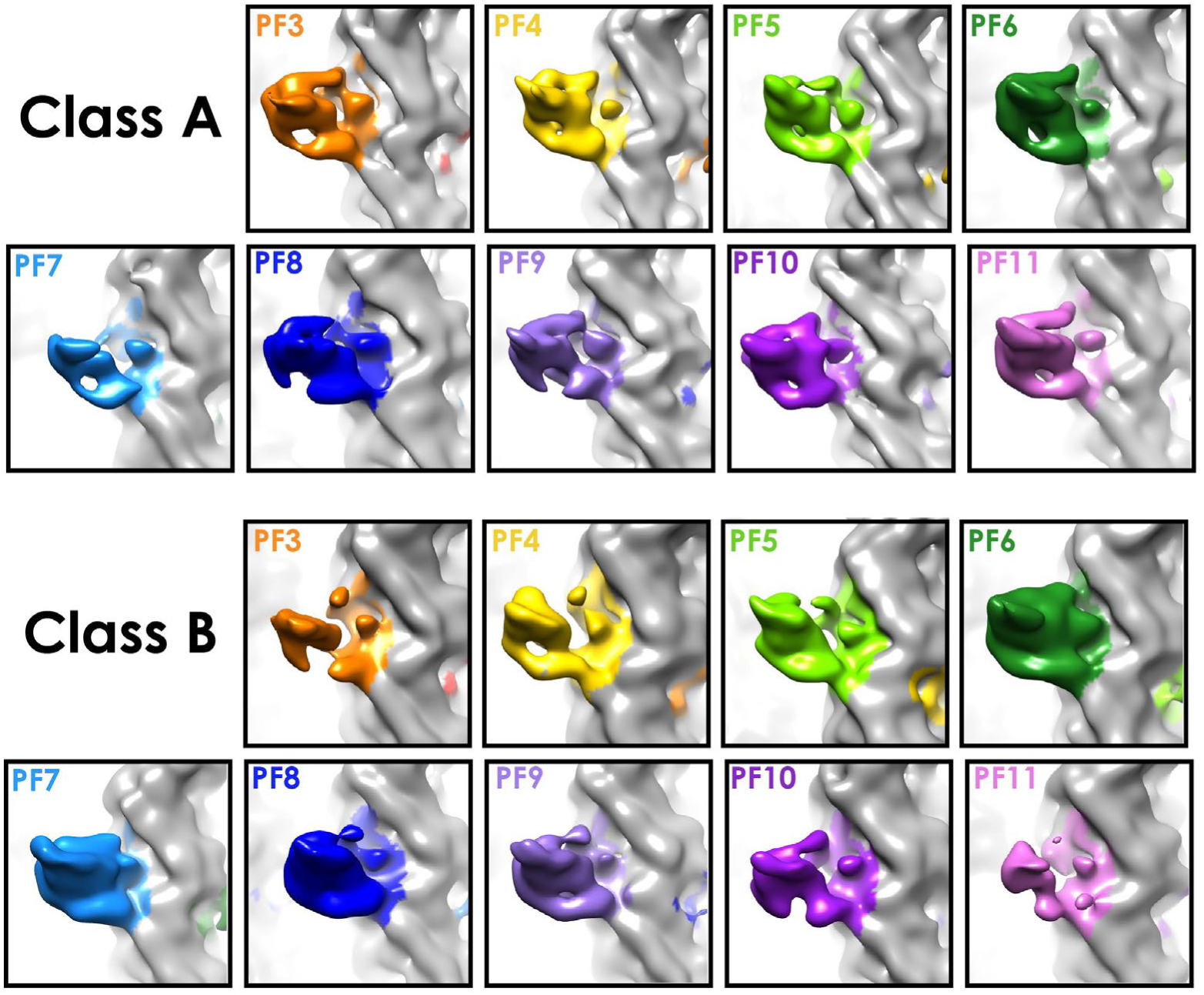
Density for each protofilament used for decoration measurements in Figure 4D and 4E.

## Movie 1

A movie depicting a morph between Classes A and B, depicting their cross-sectional distortion

